# A genetically encoded selection for amyloid-β oligomer binders

**DOI:** 10.1101/2025.03.11.642651

**Authors:** ByungUk Lee, John A. Mannone, Tina Wang

## Abstract

Soluble amyloid beta oligomers (AβOs) are a hypothesized source of neurotoxicity in Alzheimer’s Disease. Binding proteins that recognize these species may have high utility in diagnostic and therapeutic applications. However, identifying binders that recognize AβOs directly generated from the aggregation cascade is made challenging by the short lifetime and low concentrations of oligomer populations. We report a new strategy for detecting binding to AβOs as they form during Aβ42 aggregation using a genetically encoded biosensor. We show that our method enables rapid and highly reproducible measurement of the activity of existing AβO binders and can be used to select for new binders with improved potency. We uncover hits that are >20 fold more effective than reported binders at delaying secondary nucleation, the step in Aβ aggregation thought to generate the highest amounts of toxic oligomers. Our approach may greatly accelerate the discovery and characterization of binding proteins that target AβOs.

## Introduction

Alzheimer’s Disease (AD) is an aging-linked neurodegenerative disorder affecting millions worldwide with no cure to date^1^. AD pathology is linked to the misfolding of two proteins, amyloid beta (Aβ) and tau, whose presence in insoluble aggregates in the brains of affected patients is a hallmark of disease^2–4^. Foundational biophysical studies have established that these aggregates form through a cascade of self-association events involving multiple structurally distinct species including monomers, oligomers, protofibrils and mature amyloid fibrils (**Fig. 1a**)^5^. Additionally, fibril surfaces can catalyze further formation of oligomers and fibrils through a secondary nucleation mechanism, a process hypothesized to contribute to the disease progression^6–8^. While the molecular mechanism(s) responsible for AD remains to be conclusively identified, soluble Aβ oligomeric intermediates (AβOs) formed in the aggregation cascade have been implicated as a major driver of neuronal cell toxicity and death^9–11^.

**Figure 1.**
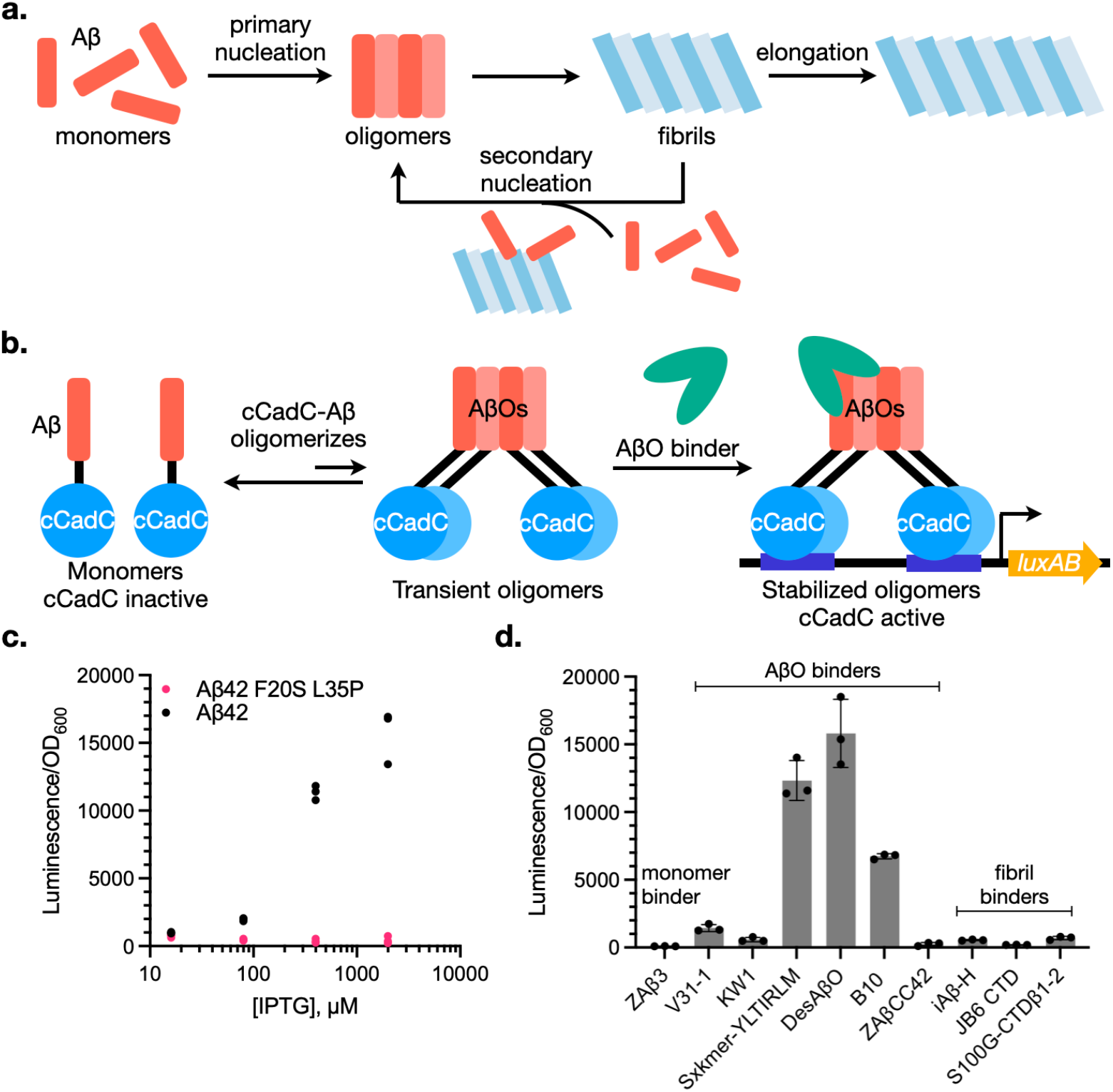
Biosensor strategy for detecting Aβ oligomer binding activity. (**a**) Overview of the Aβ42 aggregation cascade, consisting of the microscopic steps of primary nucleation of monomers to oligomeric and fibrillar species, elongation of fibril ends by monomers, and secondary nucleation from fibril surfaces, which catalyze the conversion of monomers to oligomers^6–8^. (**b**) cCadC-Aβ42 fusions link the binding and stabilization of Aβ oligomers to reporter gene transcription. (**c**) Effect of Sxkmer-YLTIRLM co-expression on *luxAB* transcriptional activation by cCadC-Aβ42 or cCadC-Aβ42 F20S L35P monomeric mutant. Data shows three biological replicates plotted individually. (**d**) Activation of cCadC-Aβ42 biosensor by reported binders of Aβ42 monomers (ZAβ3), oligomers (V31-1, KW1, Sxkmer, DesAβO, B10, and ZaβCC42), and fibrils (IAβ-H, JB6 CTD, and S100G-CTDβ1-2)^15,16,20,32,43–47^. All binders are induced using 1 mM IPTG. See **Supplementary Fig. 1** for full dose-response data. Data reflects mean and standard deviation (s.d.) of three biological replicates.

Accordingly, binding proteins, such as antibodies and antibody mimetics, that target these oligomeric species have attracted significant interest due to their potential utility in diagnostic and therapeutic applications^12,13^.

The identification of binders that recognize AβOs generated through the aggregation cascade is a challenging endeavor^12,14–16^. Because oligomers are highly transient and formed at significantly lower concentrations compared to other Aβ species^17^, immunization with native Aβ is unlikely to generate oligomer-specific antibodies. Screening or selecting for binders *in vitro* is difficult for similar reasons. The majority of reported binders have been identified using AβOs that are artificially stabilized through strategies that include covalent linkage^18–21^, chemical treatment^22–32^, and installing point mutations that promote oligomerization^33^, which can then be employed as epitopes in immunization or phage display. AβO binders have also been generated by immunizing with Aβ regions thought to be exposed in oligomers^13,34,35^ and through grafting sequences rationally designed to interact with Aβ, followed by screening for binding to immobilized oligomers^16,36,37^.

Although the binders generated through such methods have served as valuable research tools and diagnostic reagents, oligomer stabilization strategies for AβO binder discovery may potentially generate non-native epitopes, resulting in lower affinity for native AβOs generated by aggregation pathways in biological systems^12,38^. Many oligomer binders also recognize other Aβ species, such as monomers or fibrils^14,39^. Cross-reactivity can convolute a binder’s mechanism of action and decrease efficacy^13^, while in therapeutic settings strong fibril targeting is often associated with amyloid-imaging related abnormalities (ARIA-E)^39–41^. Reliable quantification of AβO concentrations for diagnostic purposes also requires high affinity and selectivity for oligomers^42^. Unfortunately, selecting for and optimizing binding to natively generated AβOs is difficult due to the challenges in assessing such activity. Existing methods employ *in vitro* biophysical measurements that are technically demanding, time-intensive, and costly. A higher-throughput approach for measuring binding to AβOs as they are generated through Aβ aggregation could greatly aid in our ability to generate oligomer binders with improved activity.

Here, we report the development of a genetically encoded biosensor that detects the ability of different proteins to bind AβOs as they form through Aβ42 aggregation in an *E. coli* model system. Our approach allows measurement of binder activity against non-stabilized oligomers and can be used to rapidly identify critical residues for activity in existing AβO binders. We apply this biosensor towards selecting for new AβO binders, resulting in the identification of an improved binder that recognizes oligomers generated from Aβ42 secondary nucleation and is > 20-fold more effective at inhibiting this step of the aggregation cascade compared to reported AβO binders. Interestingly, additional efforts to select for binders active at lower stoichiometries to Aβ led to the discovery of highly active hits that exhibit oligomerization behavior reminiscent of some ATP-independent molecular chaperones. Finally, we perform a head-to-head comparison of the activity of several AβO binders on a panel of Aβ42 mutants to characterize interactions potentially important for oligomer recognition. We envision that our method will serve as a useful complement to existing biophysical techniques for AβO binder characterization and may provide a new avenue for the discovery of potent oligomer binders with therapeutic and diagnostic utility.

## Results

### Development of a genetically encoded biosensor to measure AβO binder activity

Our overall strategy for detecting AβO binding is shown in **Fig. 1b**. Here, we employ an engineered oligomerization-dependent transcription factor, cCadC^48^, which is monomeric and inactive on its own but is activated when self-associated through fusion to oligomerizing proteins to drive transcription of its regulated promoter P_*cadBA*._ Fusing Aβ42 to cCadC could theoretically turn on cCadC transcription upon AβO formation. However, because AβOs are highly transient, we hypothesized that stabilization by a co-expressed AβO binder would be required to give the oligomeric cCadC-Aβ42 fusion sufficient time to activate gene transcription before dissociating to monomers or further aggregating into amyloid fibrils (**Fig. 1b**). Thus, this approach would enable us to measure AβO binder activity as a function of cCadC-regulated expression of reporter genes that generate easily detectable outputs (ex. luciferase).

Several considerations led to our decision to use an *E. coli* cellular model for measuring AβO binder activity. Aβ aggregation in *E. coli* has been well-studied and is highly predictable, and the *E. coli* intracellular environment should buffer Aβ from external factors that can lead to experimental artifacts and irreproducibility in *in vitro* aggregation experiments^49,50^. Like Aβ fibrils generated *in vitro*, Aβ aggregated into *E. coli* inclusion bodies also bind the amyloid-specific dye thioflavin T (ThT) and can seed Aβ monomer aggregation ^51–54^. Additionally, Aβ aggregation inhibitors identified in *E. coli* models exhibit activity both in *in vitro* assays with purified Aβ peptide and in neuronal cultures^55–57^. These observations suggest that Aβ expressed in *E. coli* assumes conformations found in physiologically relevant biological model systems.

To test our approach, we tested if Sxkmer-YLTIRLM, a recently discovered binder that recognizes AβOs generated from secondary nucleation (**Fig. 1b**)^15^, could activate bacterial luciferase (*luxAB*) transcription by cCadC-Aβ42. Indeed, induction of Sxkmer-YLTIRLM expression increased *luxAB* transcription in an inducer concentration-dependent manner (**Fig. 1c** and **Supplementary Fig. 1**), reflecting stabilization of cCadC-Aβ42 oligomers through Sxkmer-YLTIRLM binding. In contrast, in the absence of an AβO binder, cCadC-Aβ42 is transcriptionally inactive, likely reflecting its aggregation into inclusion bodies (**Supplementary Fig. 1**). cCadC activity was dependent on Aβ42 aggregation propensity, as we observed no luminescence when the monomeric Aβ42 F19S/L34P mutant^58^ was employed instead (**Fig. 1b**). This suggests that Sxkmer-YLTIRLM is binding to AβOs natively formed in *E. coli* rather than forcing Aβ42 monomers to artificially oligomerize. Finally, co-expression of binders that recognize Aβ species other than oligomers (ex. monomers or fibrils) (**Supplementary Table 1**) failed to activate cCadC-Aβ42 transcription (**Fig. 1d** and **Supplementary Fig. 2**).

We also tested additional binders that have been hypothesized to recognize specific AβO species in our assay (**Fig. 1d, Supplementary Figs. 2** and **3**, and **Supplementary Table 1**). DesAβO, a rationally designed nanobody that recognizes AβOs formed in AD mouse models, activated luciferase expression to similar levels as Sxkmer-YLTIRLM. B10, a nanobody that stabilizes protofibrils and stains Aβ plaques in hippocampal sections from AD patients, also yielded luminescence, albeit to a lesser extent. In contrast, nanobodies KW1 and V31-1 did not significantly activate *luxAB* transcription. KW1 is specific to Aβ40 oligomers and does not recognize Aβ42 oligomers, which may explain our inability to detect its binding activity. V31-1 recognizes low molecular weight Aβ42 oligomers and stains intraneural Aβ inclusions. However, V31-1 was also reported to bind Aβ monomers. Consistent with this, we observed that both V31-1 and ZAβ3 increased the solubility of a Aβ42-GFP reporter when co-expressed in *E. coli*, while Sxkmer-YLTIRLM and B10 did not (**Supplementary Fig. 4**). Therefore, it is unclear whether our inability to detect V31-1 activity arises because cCadC-Aβ42 does not populate the specific AβO species V31-1 recognizes, or because stabilization of Aβ monomers by V31-1 leads to inhibition of cCadC (which is inactive when monomeric). Taken together, these results show that our method can detect the activity of a variety of AβO binders, including those that recognize AβO conformations found in disease models.

### Identifying the sequence determinants of AβO binder activity

We tested if our method could be used to identify the features of AβO binders that are most critical for AβO binding activity, thus offering an alternative to existing lower-throughput and more technically demanding approaches for assessing AβO binding activity. Installing a R39A or R61A mutation in the complementarity-determining region (CDR) loops of B10 decreases B10 binding affinity to Aβ by ∼6- and ∼1.5-fold, respectively (**Supplementary Table 2**)^59^. We measured the effects of these mutations on B10’s ability to activate cCadC-Aβ42 transcription in our system and observed a decrease in luminescence signal upon the elimination of these positively charged residues, consistent with their effects on B10 binding affinity, suggesting that the cCadC-Aβ42 biosensor is sensitive to changes in AβO binder activity (**Fig. 2a**).

**Figure 2.**
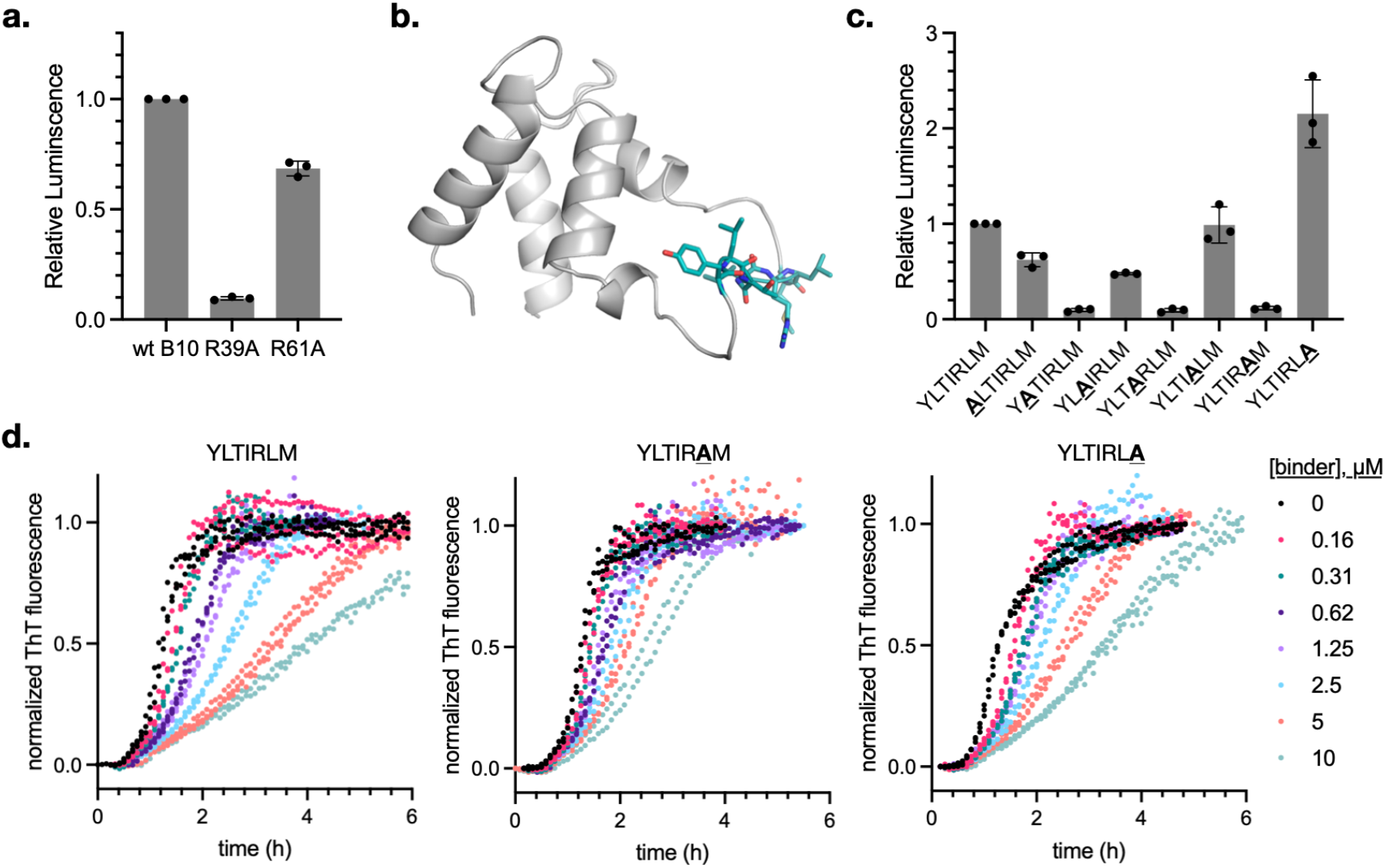
Characterization of the contribution of AβO binder features to activity. (**a**) Effect of CDR loop mutations on B10 recognition of cCadC-Aβ42 oligomers. All binders are induced using 1 mM IPTG. Data shows the luminescence signal of each mutant normalized by the luminescence signal generated by the parent B10 sequence and reflects the mean and s.d. of three biological replicates. Raw luminescence data is provided in **Supplementary Fig. 5a**. (**b**) AlphaFold2 predicted structure of Sxkmer-YLTIRLM, with the loop sequence colored in teal. (**c**) Effect of single alanine mutations of the loop sequence on Sxkmer-YLTIRLM recognition of cCadC-Aβ42 oligomers. All binders are induced using 1 mM IPTG. Data shows the luminescence signal of each alanine mutant normalized by the luminescence signal generated by the parent YLTIRLM sequence and reflects the mean and s.d. of three biological replicates. Raw luminescence data is provided in **Supplementary Fig. 5b**. (**d**) Comparison of the activities of Sxkmer-YLTIRLM, YLTIRAM, and YLTIRLA in *in vitro* ThT fluorescence assays of Aβ42 aggregation. Assay conditions: 3 µM Aβ42 in 20 mM Sodium Phosphate, 0.2 mM EDTA, pH 8.0. Data reflects three technical replicates plotted as individual values.

We also performed alanine scanning of the Sxkmer-YLTIRLM loop sequence to determine the contribution of the individual amino acids in this region to AβO binding. Sxkmers consist of a scaffold derived from the S100G calcium binding protein grafted with a 7-residue loop sequence that mediates epitope binding (**Fig. 2b**). Y48A, T50A, and R52A substitutions in the YLTIRLM loop minimally impacted binding activity, whereas mutation of L49, I51, and L53 to alanine greatly decreased cCadC-Aβ42 oligomer binding (**Fig. 2c**), suggesting that bulky aliphatic sidechains at these positions may play a prominent role in AβO recognition.

Surprisingly, the M54A mutation increased luminescence signal, potentially suggesting an increase in binding activity or binder expression level for this mutant (**Fig. 2c**). To determine if observations from the cCadC-Aβ42 biosensor translate into binder activity *in vitro*, we compared the effects of purified Sxkmer-YLTIRLM, YLTIRAM and YLTIRLA (**Supplementary Fig. 6**) on Aβ42 aggregation kinetics in a ThT fluorescence assay (**Fig. 2d**). These experiments revealed a large decrease in Sxkmer-YLTIRAM’s ability to inhibit aggregation compared to Sxkmer-YLTIRLM, suggesting a loss of Aβ42 recognition, while in contrast, Sxkmer-YLTIRLA showed activity comparable to Sxkmer-YLTIRLM. The discrepancy between Sxkmer-YLTIRLA activities measured *in vitro* and through our method may arise from differences in binder expression or differences in sensitivity to small changes in binder activity between the two assays. Together, these results suggest that our method can be used to rapidly identify binder features important for AβO recognition.

### Discovery of a new AβO binder

We adapted the cCadC biosensor into a selection for identifying new AβO binders by placing chloramphenicol acetyltransferase (*cat*), which confers resistance to the antibiotic chloramphenicol, under control of P_*cadBA*_ in place of *luxAB* (**Fig. 3a**). *E. coli* cells encoding this selection must express a protein that binds cCadC-Aβ42 oligomers to survive in the presence of chloramphenicol. Indeed, cells expressing Sxkmer-YLTIRLM grew extensively on media containing 25 µg/mL chloramphenicol, while cells with the Sxkmer-YLTIRAM mutant experienced severe growth inhibition, demonstrating excellent discrimination of AβO binding activity by our selection (**Fig. 3b**).

**Figure 3.**
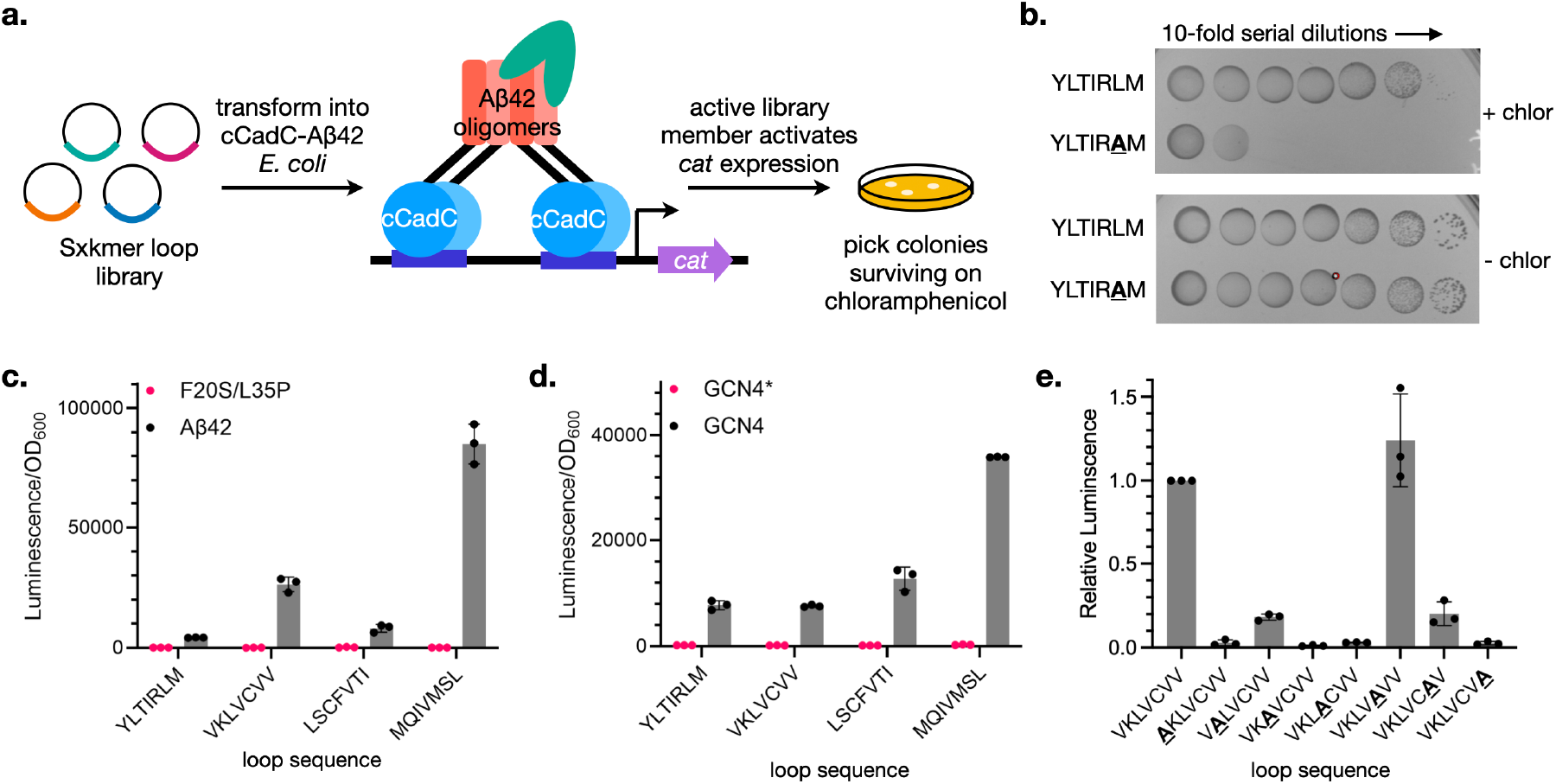
Selection for new AβO binders. (**a**) Overview of cCadC-Aβ42 based selection strategy for new AβO binders from a Sxkmer loop library. (**b**) Growth of selection cells encoding Sxkmer-YLTIRLM or activity-deficient mutant Sxkmer-YLTIRAM on chloramphenicol-containing media (top) or media containing maintenance antibiotics only (bottom). (**c**) Activation of cCadC fused to Aβ42 or monomeric mutant F20S L35P by identified hits. (**d**) Activation of cCadC fused to an unrelated monomeric (GCN4*) or dimeric (GCN4) peptide by identified hits. (**e**) Alanine scanning analysis of Sxkmer-VKLVCVV loop sequence. Data shows the luminescence signal of each alanine mutant normalized by the luminescence signal generated by the parent VKLVCVV sequence. Raw luminescence data is provided in **Supplementary Fig. 5c**. Binders in (**c**-**e**) were induced using 1 mM IPTG. Data reflects mean and s.d. of three biological replicates.

Next, we generated a library of Sxkmers where the 7 residues in the loop sequence were fully randomized using site-saturation mutagenesis (library size ∼10^9^) and subjected cells harboring this library (∼10^8^ clones) to selection for members with AβO binding activity. Cells were first plated on solid media containing 40 µg/mL chloramphenicol and 1 mM IPTG to induce Sxkmer production. Because cells expressing Sxkmer-YLTIRLM exhibited minimal growth on media with this chloramphenicol concentration, we hypothesized that these conditions might enable us to identify more active AβO binders. Library members from ∼200 surviving colonies were cloned back into the parent vector and subjected to a second round of selection using an increased chloramphenicol concentration (80 µg/mL) to identify more active variants. Sanger sequencing of the eight surviving colonies after this second round revealed three distinct loop sequences, VKLVCVV, LCSFVTI, and MQIVMSL.

All three newly identified Sxkmer variants displayed a greater degree of cCadC-Aβ42 transcriptional activation compared to Sxkmer-YLTIRLM, potentially reflecting an increased ability to bind AβOs (**Fig. 3c**). However, we also anticipated several ways in which selected Sxkmers might “cheat” our selection and tested for each scenario. All Sxkmer variants failed to induce *luxAB* transcription when tested with the cCadC-Aβ42 F20S/L35P mutant, confirming that they do not artificially force Aβ42 oligomerization (**Fig. 3c**). Furthermore, none of the hits activated cCadC fused to an unrelated monomeric protein (GCN4*)^48^, suggesting that they are also not inducing self-association of the cCadC protein itself (**Fig. 3d**). However, Sxkmer-LCSFVTI and Sxkmer-MQIVMSL enhanced *luxAB* transcription by cCadC fused to an unrelated dimeric protein (GCN4)^48^, revealing that these hits are false positives that bind to and stabilize cCadC that is already self-associated (**Fig. 3d**). In contrast, Sxkmer-VKLVCVV failed to enhance cCadC-GCN4 transcriptional activity, suggesting that this is a true positive that specifically binds to AβOs (**Fig. 3d**). Alanine scanning of the VKLVCVV loop sequence revealed that both the bulky aliphatic and positively charged residues are key contributors to activity (**Fig. 3e**). Only mutation of the cysteine did not lead to reduced cCadC-Aβ42 activation, suggesting that this position may be amenable to further derivatization (**Fig. 3e**).

We next determined whether the large increases in cCadC-Aβ42 activation by Sxkmer-VKLVCVV compared to Sxkmer-YLTIRLM translated into increased potency *in vitro*. We purified both binders (**Supplementary Fig. 6**) and tested their activity in ThT fluorescence assays of Aβ42 aggregation (**Fig. 4a**). Both Sxkmer-YLTIRLM and VKLVCVV delayed Aβ42 aggregation, primarily exerting their effects on the slope of fibril formation. The aggregation kinetics fit well with a model where the secondary nucleation step is selectively inhibited (**Fig. 4a** and **Supplementary Fig. 7**). This suggests that, like Sxkmer-YLTIRLM, VKLVCVV may delay Aβ42 aggregation through binding oligomers generated from secondary nucleation. Moreover, Sxkmer-VKLVCVV appears to be a stronger inhibitor of secondary nucleation compared to Sxkmer-YLTIRLM, enabling a > 20-fold higher reduction in the secondary nucleation rate constant (k_2_) when added at 5 µM (**Fig. 4b**). To obtain insight into Aβ42 morphologies stabilized by these binders, we examined reaction mixtures containing Aβ42 in the presence or absence of Sxkmer-YLTIRLM or VKLVCVV after a 3.5 h incubation at 37 °C using transmission electron microscopy (TEM) (**Fig. 4c-e**). In the absence of binders, Aβ42 forms aggregates with some fibrillar morphology (**Fig. 4c**) which are eliminated in the presence of both Sxkmer-YLTIRLM (**Fig. 4d**) and VKLVCVV (**Fig. 4e**).

**Figure 4.**
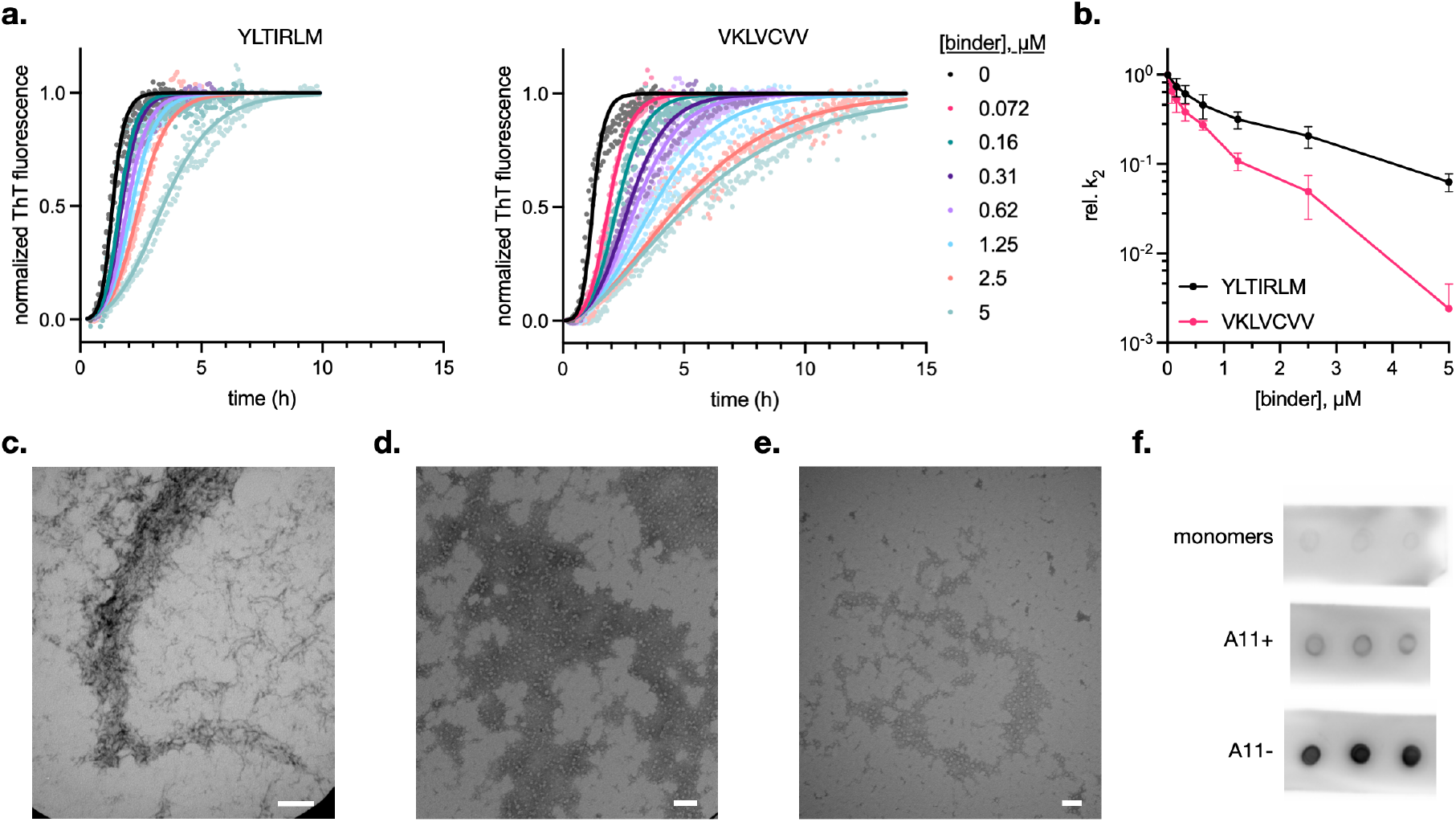
*In vitro* characterization of Sxkmer-VKLVCVV activity. (**a**) Comparison of the activities of Sxkmer-YLTIRLM and VKLVCVV in *in vitro* ThT fluorescence assays of Aβ42 aggregation. Assay conditions: 3 µM Aβ42 in 20 mM Sodium Phosphate, 0.2 mM EDTA, pH 8.0. Data reflects three technical replicates plotted as individual values. Lines represent fits to a selective reduction of the secondary nucleation rate constant using the AmyloFit software^60^. (**b**) Effect of Sxkmer-YLTIRLM and VKLVCVV on the secondary nucleation rate constant (k_2_) of Aβ42 aggregation. Data represent mean and s.d. of four separate replicates of ThT fluorescence assay data except for the 5 µM VKLVCVV data point, which represents the mean and s.d. of two replicates. (**c**-**e**) Transmission electron microscopy (TEM) analysis of 3 µM Aβ42 incubated either alone (**c**) or in the presence of 3 µM Sxkmer-YLTIRLM (**d**) or Sxkmer-VKLVCVV (**e**) for 150 mins under quiescent conditions. (**f**) Binding of Sxkmer-YLTIRLM, YLTIRAM, or VKLVCVV to two different AβO species generated through chemical treatment. A11+ oligomers are spherical, non-fibrillar oligomers stained by the A11 antibody^61,62^. A11-oligomers are not recognized by the A11 antibody and represent larger oligomeric species.

Finally, we conducted experiments to further clarify Sxkmer-VKLVCVV’s mechanism of action. We first measured Sxkmer-VKLVCVV affinity to immobilized Aβ42 fibrils by SPR to determine whether its observed activity in ThT assays might instead stem from significant binding to fibril surfaces that catalyze secondary nucleation. Sxkmer-VKLVCVV displayed weak binding to fibrils, with an apparent K_d_ of ∼ 50 µM (**Supplementary Fig 8**), well above concentrations at which it exhibits significant inhibitory activity in ThT assays (**Fig. 4a-b**). This suggests that Sxkmer-VKLVCVV does not primarily inhibit secondary nucleation through binding fibril surfaces. We also used dot blotting to assess the binding of Sxkmer-VKLVCVV sequences to Aβ42 monomers and two different AβO species generated by chemical treatment according to reported protocols^61,62^. Sxkmer-VKLVCVV appears to recognize both chemically stabilized AβO species, but stains A11-oligomers more strongly, and furthermore exhibits negligible recognition of monomers (**Fig. 4f**). Together, these results demonstrate that the cCadC-Aβ42 biosensor can be employed to identify new AβO binders with an increased ability to inhibit the secondary nucleation step of Aβ42 aggregation.

### Selection for highly active AβO binders

We next determined if we could identify Aβ oligomer binders that are highly active even when employed at lower concentrations by repeating the above selection using a 100-fold lower expression level of the Sxkmer library. Cells were transformed with the same Sxkmer loop library used to identify Sxkmer-VKLVCVV and plated on solid media containing 40 µg/mL chloramphenicol and 0.01 mM IPTG to induce Sxkmer production. Loop sequences from 48 surviving colonies were subcloned back into the parent vector and subjected to a second round of selection using a higher chloramphenicol concentration (50 µg/mL) while maintaining Sxkmer induction at 0.01 mM IPTG. Sequencing of the 8 surviving colonies revealed enrichment of three new loop sequences: IYILVER, FRFFIYF, and YRFFIYW (**Fig. 5a**). In the case of IYILVER, we also identified a premature stop codon in the Sxkmer S100G scaffold that deletes the C-terminus of the S100G protein starting partway through helix III (**Fig. 5b**). Moving forward, we refer to this truncated scaffold as “tSxkmer.”

**Figure 5.**
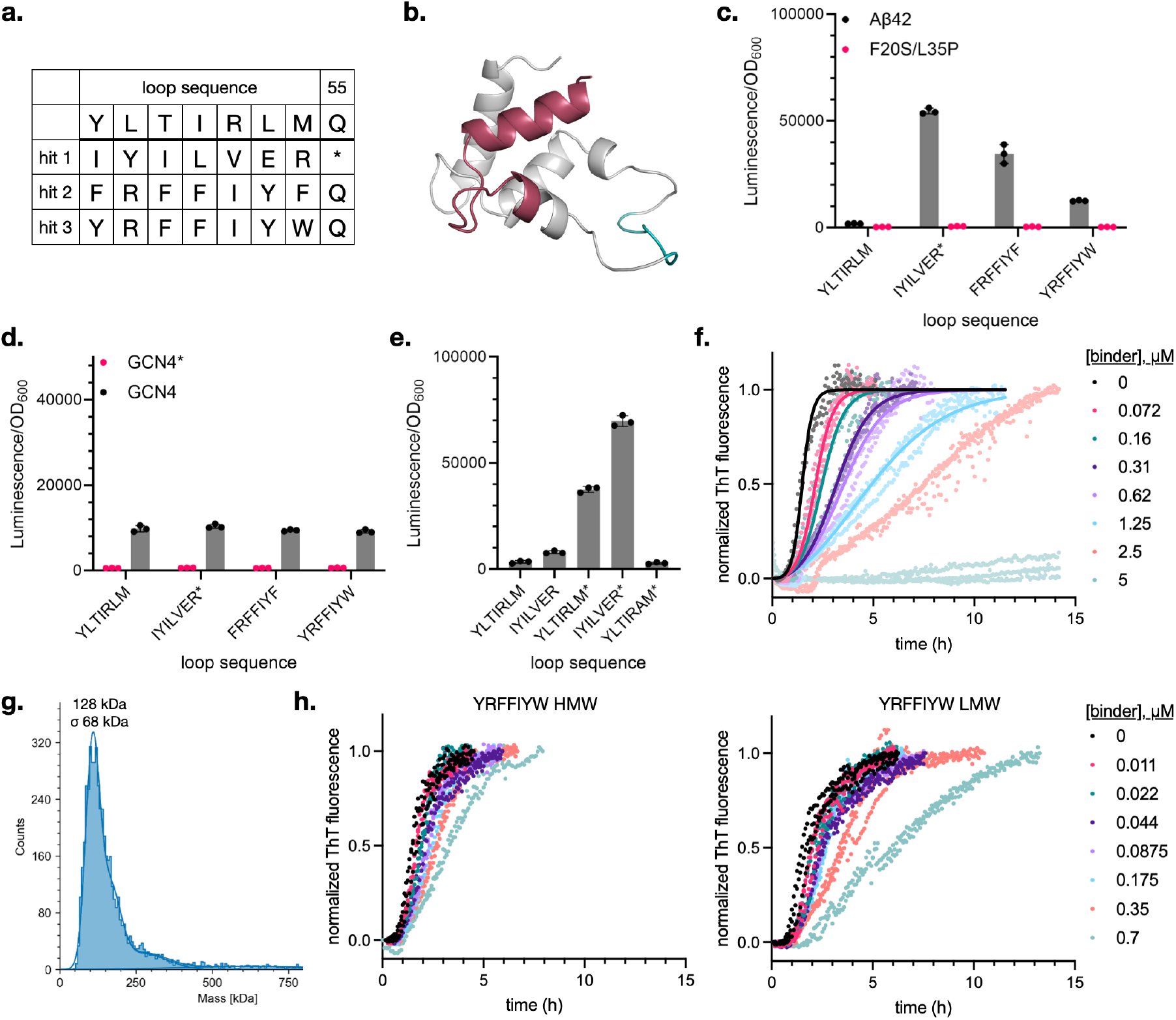
Identification of Aβ oligomer binding Sxkmer loop sequences with activity at lower binder concentrations. (**a**) Loop sequences of hits obtained from selections using a lower expression level of Sxkmer library members. Hit 1 also contains a mutation in the S100G scaffold sequence (Q55*; refers to S100G residue numbering). (**b**) AlphaFold2 structure of the Sxkmer scaffold with the region truncated by Q55* shown in red. The loop sequence is colored in teal. (**c**) Activation of cCadC fused to Aβ42 or monomeric mutant F20S L35P by identified hits. Asterisk indicates a tSxkmer scaffold. (**d**) Activation of cCadC fused to an unrelated monomeric (GCN4*) or dimeric (GCN4) peptide by identified hits. Asterisk indicates a tSxkmer scaffold. (**e**) Effect of the scaffold truncation on the activity of Sxkmer loop sequences. Asterisk indicates a tSxkmer scaffold. Binders in (**c**-**e**) were induced using 0.01 mM IPTG. (**f**) Activity of Sxkmer-YRFFIYW in *in vitro* ThT fluorescence assays of Aβ42 aggregation. Assay conditions: 3 µM Aβ42 in 20 mM Sodium Phosphate, 0.2 mM EDTA, pH 8.0. (**g**) Mass photometry analysis of Sxkmer-YRFFIYW oligomers. Average molecular weight of the Sxkmer-YRFFIYW monomer (including 6xHis tag) is 11250 Da. (**h**) Comparison of the activities of high molecular weight (HMW) and low molecular weight (LMW) Sxkmer-YRFFIYW fractions from SEC (see **Supplementary Fig. 15** for SEC trace) in *in vitro* ThT fluorescence assays of Aβ42 aggregation. Assay conditions: 3 µM Aβ42 in 20 mM sodium phosphate, 0.2 mM EDTA, pH 8.0. Binder concentrations are calculated using monomeric Sxkmer-YRFFIYW molecular weight. (**c**-**e**) Data reflects mean and s.d. of three biological replicates. (**f, h**) Data reflects three technical replicates plotted as individual values.

All three hits were more active than Sxkmer-YLTIRLM or VKLVCVV in activating cCadC-Aβ42 transcription, with luminescence signal developing at much lower expression levels (**Fig. 5c** and **Supplementary Fig. 9**). Unexpectedly, higher induction of these new hits led to reduced luminescence signal, which was not observed in the case of Sxkmer-YLTIRLM or VKLVCVV. None of the new hits activated cCadC-Aβ42 F20S/L35P or cCadC-GCN4*, demonstrating that they do not force oligomerization of Aβ42 or cCadC (**Fig. 5d**). Furthermore, the new hits did not magnify transcriptional activity of cCadC-GCN4 (**Fig. 5d**), except when Sxkmer-FRFFIYF and YRFFIYW were induced using high IPTG levels (**Supplementary Fig. 10**). This suggests that, at low binder concentrations, these hits are selective for Aβ42, but may begin to exhibit nonspecific activity when used at higher levels.

C-terminal truncation of the Sxkmer scaffold in the hit with loop sequence IYILVER was an unexpected outcome of our selection, and we elected to characterize its effect on Sxkmer activity. When placed back into a full-length Sxkmer scaffold, IYILVER induced ∼2-fold greater activation of cCadC-Aβ42 compared to Sxkmer-YLITRLM but showed ∼9-fold decreased activity relative to tSxkmer-IYILVER (**Fig. 5e**). Therefore, while the IYILVER loop sequence binds to AβOs, the truncated S100G scaffold plays a major role in the observed activity. Similarly, when we grafted the YLITRLM loop sequence onto tSxkmer, the resultant binder induced a much larger increase in cCadC-Aβ42 activation at low expression levels compared to Sxkmer-YLTIRLM (**Fig. 5e** and **Supplementary Fig. 11**). Finally, the YLTIRAM sequence, which exhibits diminished AβO binding (**Fig. 2c-d**), activates cCadC-Aβ42 transcription to a comparable degree as Sxkmer-YLTIRLM when grafted onto the tSxkmer scaffold (**Fig. 5e**).

Together, these results suggest that the tSxkmer scaffold amplifies cCadC-Aβ42 oligomer binding activity. Unfortunately, repeated efforts to purify tSxkmers were unsuccessful and precluded further characterization *in vitro*.

Next, we further characterized the non-truncated hits, Sxkmer-FRFFIYF and YRFFIYW, which share highly similar loop sequences. Alanine scanning analysis indicates that, for both hits, all seven loop residues are necessary for activity at low binder expression levels (**Supplementary Fig. 12**). Next, we purified Sxkmer-YRFFIYW (**Supplementary Fig. 5**) and tested its activity in ThT assays of Aβ42 aggregation. Sxkmer-YRFFIYW was a significantly more potent inhibitor of Aβ aggregation compared to Sxkmer-YLTIRLM when employed at lower concentrations (**Fig. 5f**). Furthermore, at higher concentrations, Sxkmer-YRFFIYW completely suppressed aggregation within the time period examined (**Fig. 5f**). Interestingly, unlike Sxkmer-YLTIRLM and VKLVCVV, which are primarily monomeric, Sxkmer-YRFFIYW forms polydisperse oligomers exhibiting molecular weight > 10mer species (**Fig. 5g** and **Supplementary Fig. 13**). Circular dichroism shows that Sxkmer-YRFFIYW retains α-helical character, indicating that the S100G scaffold is folded within the oligomeric assembly (**Supplementary Fig. 14**). To characterize the contribution of different Sxkmer-YRFFIYW oligomeric states to its activity, we resolved both a higher- and lower-molecular weight complex by SEC (**Supplementary Fig. 15**) and tested the activity of each population in ThT assays.

Interestingly, the lower-molecular weight (LMW) population was significantly more active in inhibiting Aβ42 aggregation compared to the high-molecular weight (HMW) population (**Fig. 5h**).

Kinetic fitting of the ThT assay data suggests that Sxkmer-YRFFIYW is a secondary nucleation inhibitor (**Fig. 5f** and **Supplementary Fig. 16**), similar to Sxkmer-YLTIRLM and VKLVCVV. SPR analysis of Sxkmer-YRFFIYW binding to immobilized Aβ42 fibrils indicated that it has a higher affinity (K_d_ ∼ 4.7 µM) for fibrils compared to Sxkmer-VKLVCVV (**Supplementary Fig 17**). At lower concentrations, Sxkmer-YRFFIYW likely suppresses secondary nucleation through binding AβOs, but at higher concentrations it may begin to recognize fibrils. Thus, the complete suppression of Aβ42 aggregation observed when Sxkmer-YRFFIYW is added at 5 µM in ThT assays may stem from a different mechanism involving fibril binding (**Fig. 5f**).

### Characterization of AβO:binder interactions using a Aβ42 single point mutant panel

We determined the activity of both reported AβO binders and our newly identified Sxkmer variants on a panel of cCadC-Aβ42 single point mutants to characterize Aβ42 residues important for binder:AβO interactions (**Fig. 6**). Mutations in Aβ may affect oligomerization propensity, oligomer heterogeneity, and/or recognition by AβO binders. Although this pleiotropy convolutes our experiments, we hypothesized that AβO binders may utilize distinct binding mechanisms that would enable us to identify general patterns of cCadC activation with different Aβ42 mutants. Additionally, we included Sxkmer-MQIVMSL, which stabilizes self-associated cCadC (**Fig. 3d**), which should in principle control for the effect of mutations on global Aβ42 oligomerization. In addition to Aβ42 mutations linked to familial Alzheimer’s Disease^63^, we selected mutations that should induce a large change in side-chain properties but exert minimal effects on nucleation propensity^64,65^.

We tested over 100 combinations of binders and Aβ42 mutants that exhibited a wide range of transcriptional activation (**Fig. 6**). The inter-day reproducibility of binder activity on these mutants was excellent (R^2^ > 0.95; **Supplementary Fig. 19**), highlighting the robustness of the cCadC-Aβ42 assay. Generally, Aβ42 mutants reported to increase oligomerization or aggregation propensity (E22G, E22Δ, D23N, G33A, and Aβ43) led to increases in cCadC transcription across all binders. The exception was E22G, which decreased luminescence signal from the new Sxkmer variants identified in the highest stringency selection (tSxkmer-IYILVER, Sxkmer-FRFFIYF, and Sxkmer-YRFFIYW). Otherwise, all new Sxkmers arising from the cCadC selections exhibited a similar pattern of activity across the Aβ42 mutants. The S100G scaffold itself may be biasing oligomer binding, which is consistent with the observation that Sxkmer-YLTIRLM and the cCadC binder Sxkmer-MQIVMSL partially share the same activity pattern. B10 and DesAβO exhibit a somewhat different activity profile on the Aβ42 mutant panel. The Aβ42 R5G mutation has a particularly bifurcated effect on AβO binders depending on the scaffold, indicating that R5 contributes to the interactions of Sxkmer-based oligomer binders but not nanobodies. Sxkmer-YLTIRLM’s activity greatly increased upon loss of negative charges at E22 and D23, while both the newly identified Sxkmer variants and the nanobodies demonstrated a more muted response. Because all Sxkmer-derived AβO binders we tested inhibit secondary nucleation, this could indicate that Sxkmer-YLITRLM and the new Sxkmer variants are using different interactions to bind the same cCadC-AβO species generated by secondary nucleation. Another point of difference between Sxkmer-YLTIRLM and the newly selected Sxkmers is their activity on the V18N mutant. V18N greatly diminishes activity of the new Sxkmer variants, suggesting that interaction with V18 may be critical for their recognition of AβOs.

**Figure 6.**
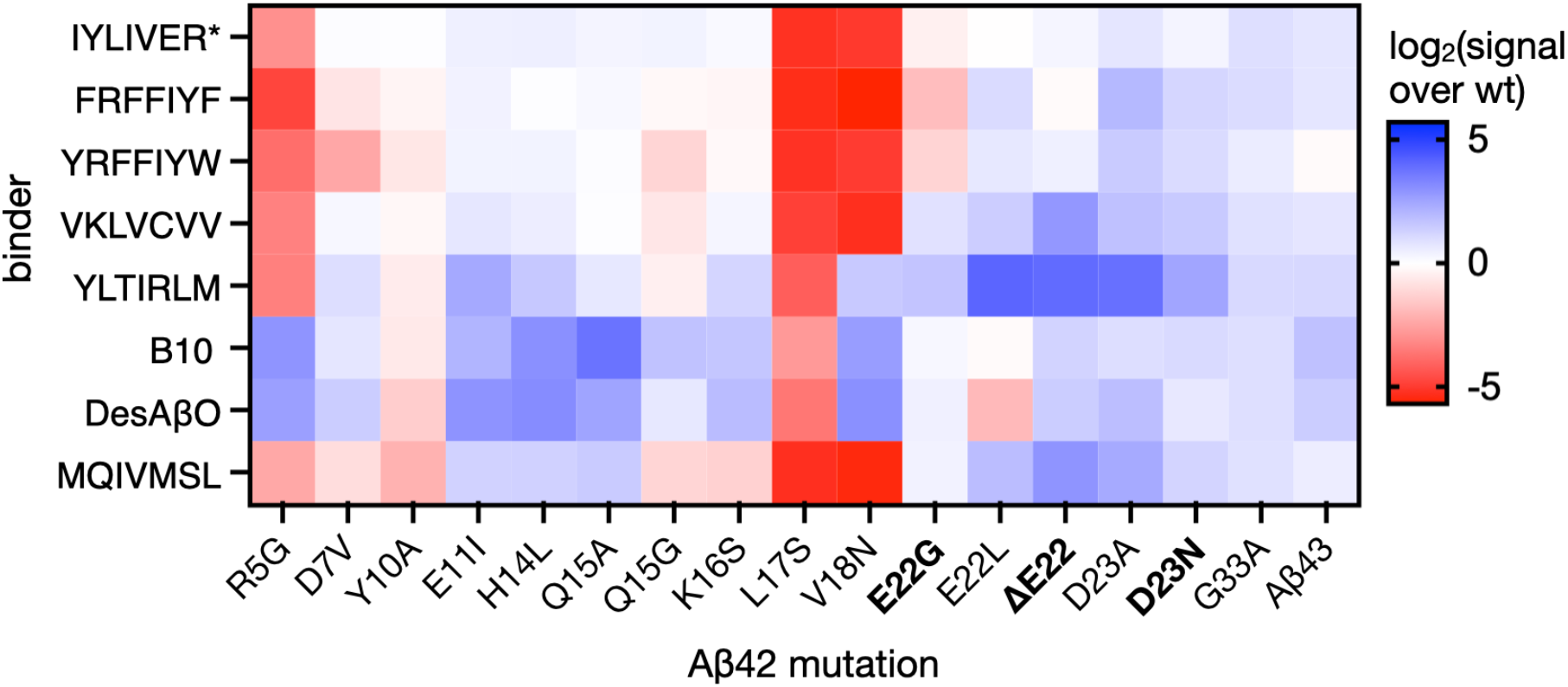
Profiling of binder activity on cCadC-Aβ42 mutants. Effect of mutations in the Aβ sequence on cCadC-Aβ42 transcriptional activation by AβO binders examined in this study. Bolded mutations are familial Alzheimer’s Disease associated. Data corresponds to the log_2_ of luminescence signal from Aβ42 mutants normalized against the signal from the WT Aβ42 sequence for each binder. Data reflect the mean of biological duplicates. Raw luminescence data is provided in **Supplementary Fig. 18**.

## Discussion

We have developed a new, scalable method for measuring the activity of AβO binders using a genetically encoded biosensor for AβO binding in an *E. coli* model. Although this approach diverges from the *in vitro* analytical techniques historically employed by the amyloid aggregation field, it has several advantageous features. First, our assay exhibits remarkably high reproducibility. The use of the *E. coli* cellular environment for Aβ oligomer formation may buffer external factors that could perturb Aβ aggregation and increase variability. By interrogating bulk populations of *E. coli*, we essentially obtain averages of ∼10^8^ independent aggregation experiments taking place in parallel in individual cells, which likely also increases the reliability of our measurements. Second, our data suggests that we are detecting binding to oligomers as they are forming during the Aβ42 aggregation cascade. This enables us to select for binders that target these transient intermediates rather than having to use chemically stabilized oligomers, which may generate non-native species. Finally, the partitioning of Aβ fibrils to insoluble inclusion bodies in *E. coli* enables high selectivity of the cCadC-Aβ42 assay for oligomer binding activity, as proteins in inclusion bodies are transcriptionally incompetent.^49^

Using our method, we were able to identify Sxkmer loop variants that exhibited more potent suppression of Aβ aggregation compared to Sxkmer-YLTIRLM, a previously reported binder that was discovered serendipitously in an effort to identify Aβ42 monomer binders. One of our hits, Sxkmer-VKLVCVV, inhibited Aβ42 secondary nucleation with > 20-fold higher activity compared to Sxkmer-YLTIRLM and exhibited low off-target binding to monomers or fibrils, thus illustrating the benefits of our ability to directly select for AβO binding and tune selection stringency to optimize hit potency. While it is not possible to measure binding affinities to oligomers produced in an ongoing aggregation reaction, we can speculate that Sxkmer-VKLVCVV’s activity arises from an increase in binding to AβOs generated from secondary nucleation. Interestingly, our experiments suggest that our other hit, Sxkmer-YRFFIYW, also recognizes AβOs made through secondary nucleation. This may suggest that oligomers formed in the cCadC-Aβ42 assay are predominantly secondary nucleation-derived, consistent with the hypothesis that most Aβ42 oligomers are generated from secondary nucleation. However, unlike Sxkmer-VKLVCVV, at higher concentrations YRFFIYW produces a marked decrease in luminescence signal in the cCadC-Aβ42 assay and completely suppresses *in vitro* Aβ42 aggregation. This behavior cannot be explained solely through an oligomer-binding mechanism. Instead, at high concentrations Sxkmer-YRFFIYW may suppress Aβ aggregation through a mixture of inhibition mechanisms.

Our efforts to select more potent Aβ42 oligomer binders resulted in the identification of hits that are themselves oligomeric despite originating from the monomeric and thermostable S100G scaffold. However, all other known members of the S100 family of proteins form dimers or oligomeric states beyond dimers^66^. Thus, it is tempting to speculate that the S100G scaffold is evolutionarily poised to oligomerize. Oligomerization is likely driven by self-association of the highly hydrophobic YRFFIYW loop. This may prevent non-specific interactions with other *E. coli* proteins or the bacterial membrane, which could otherwise lead to the binder exerting toxic effects in cells. The growth-based selection strategy we employed may have fortuitously purged toxic loop sequences and identified binder oligomerization as a possible protective mechanism. Interestingly, Sxkmer-YRFFIYW’s behavior is also reminiscent of ATP-independent chaperones, which form poly-disperse, higher-order oligomers that exchange with lower-order oligomers or monomers. This dissociation into smaller assemblies is often required for chaperone activity.

While our experiments suggest that smaller oligomeric assemblies of Sxkmer-YRFFIYW were more active in suppressing Aβ aggregation compared to larger clusters, further investigation is required to fully elucidate this binder’s mechanism of action.

The recent success of clinical trials leading to the approval of the first Alzheimer’s Disease modifying therapies (Aducanamab and Lecanemab) has led to a renewed interest in Aβ as a target. We have demonstrated that the system reported here can identify binders of Aβ oligomers that are highly active in suppressing both fibril formation and specific microscopic steps in the aggregation cascade. Moving forward, it may be interesting to use our method to identify binders using alternative scaffolds with greater precedence for diagnostic and therapeutic utility, such as nanobodies. We hope that the cCadC-Aβ42 assay will accelerate the discovery and characterization of binders that can help elucidate and address the mechanisms of Aβ oligomer-derived pathogenesis.

## Materials and Methods

### General methods

Antibiotics (Gold Biotechnology) were used at the following working concentrations: ampicillin, 50 µg/mL; spectinomycin, 100 µg/mL; chloramphenicol, 25 µg/mL; kanamycin, 50 µg/mL; tetracycline, 10 µg/mL; streptomycin, 50 µg/mL. Inducers were diluted from stock with the following concentrations: anhydrotetracycline (aTc), 100 ug/mL; arabinose, 1 M. HyClone water (GE Healthcare Life Sciences) was used for PCR reactions and cloning. For all other experiments, water was purified using a MilliQ purification system (Millipore).

Genes were obtained from Twist Bioscience as synthesized gene fragments. Phusion U Hot Start DNA polymerase (Thermo Fisher Scientific) or Q5 polymerase (New England Biolabs) were used to generate PCR products for plasmid construction. A full list of plasmids used in this work is given in **Supplementary Table 3**. Key primer sequences are listed in **Supplementary Table 4**. A full list of reagents and equipment used in this work is given in **Supplementary Table 5**.

### Bacterial strains, culture conditions, and media

*E. coli* strain S2060 was used for general plasmid cloning and amplification and bacterial luminescence and fluorescence assays. This strain is derived from strain DH10β and has the following genotype: *endA1 recA1 galE15 galK16 nupG rpsL ΔlacIZYA araD139 Δ(ara,leu)7697 mcrA Δ(mrr-hsdRMS-mcrBC) proBA::pir116 araE201 ΔrpoZ Δflu ΔcsgABCDEFG ΔpgaC λ– F’ proA+B+ Δ(lacIZY) zzf::Tn10 lacIQ1 PN25-tetR luxCDE Ppsp(AR2) lacZ luxR Plux groESL*. NEB 10β cells were used for Sxkmer-loop library cloning. BL21 DE3 pLysS cells were used for Aβ42 and AβO binder purification. *E. coli* strain S2060 *ΔcadCBA* was generated by the Lambda Red recombineering method as previously reported. Briefly, dsDNA donor, amplified from pKD4 using primers BL0293 and BL0294, was used to delete the *cadCBA* operon from strain S1021, the F-parent of strain S2060. After removing the kanamycin resistance cassette with pCP20 and subsequent curing of pCP20, the strain was F-conjugated using strain S2060 as a donor. S2060 *ΔcadCBA* was used for AβO binder selections to reduce background P_*cadBA*_ transcription from endogenous CadC.

Bacterial cell culture conditions and plasmid transformation were performed as previously described.^48^ 2xYT medium (US Biological) was used for general bacterial culture. Davis Rich Media (US Biological) was used for luminescence assays. LB media was used for expression of proteins. Auto-induction media was used for expression of Aβ42.

### Bacterial luminescence assays and GFP fluorescence assays

Overnight cultures of cells harboring the plasmids of interest were generated by inoculating single colonies in 2xYT media containing maintenance antibiotics and shaking at 37°C for 16 h. Saturated overnight cultures were diluted 1:100 into 1 mL of fresh DRM with antibiotics into 96 deep-well plates. The cultures were then incubated at 37°C with shaking (225 rpm) until they reached OD_600_ of 0.4, whereupon cells were treated with IPTG to induce binders. After shaking the cultures for an additional 30 minutes at 37°C, aTc was added into cultures to induce cCadC-Aβ42 or Aβ42-GFP. Cells were subsequently incubated at 37°C for another 3 or 6 hours for luminescence and GFP fluorescence measurements, respectively. 100 µL of culture was then transferred into 96 well black clear-bottom plates. Luminescence, fluorescence, and absorbance measurements at 600 nm were made using a Tecan Infinite Mplex plate reader. OD normalized luminescence and fluorescence measurements were generated by dividing raw luminescence and fluorescence signal by background-subtracted OD_600_ values.

### Expression of reported binders in S2060s

Saturated overnight cultures of cells harboring plasmids encoding binders were diluted 1:100 into 200 mL of fresh 2xYT media in shake flasks. The cultures were then incubated at 37 °C with shaking (225 rpm) until they reached OD_600_ of 0.4, whereupon expression of binders was induced by addition of 1 mM IPTG. Cells were then incubated for an additional 4 hours at 37 °C and harvested by centrifugation. Cell pellets were then lysed in 20 mL of lysis buffer (10 mM Tris-HCl, 0.1 mM CaCl_2_, pH 7.5) by sonication. The lysate post-sonication was used as the total fraction. The soluble fraction was collected by centrifuging the lysate at 16,000 rcf for 10 minutes and collecting the soluble fraction.

### Library generation of Sxkmer-loop variants

Primers BL0722 and BL0723 were used to amplify pBL098. The PCR product was gel-extracted and then DpnI digested to remove template DNA. The resulting mixture was then cleaned up through PCR purification and 0.5 pmol of DNA was assembled through USER assembly with added T7 DNA ligase. The assembly reaction was then subjected to PCR purification and transformed into electrocompetent *E. coli* NEB 10β cells and allowed to recover in 2xYT supplemented with 20 mM glucose at 37 °C with shaking (225 rpm) for 1 h. An aliquot was removed to estimate transformants by dilution plating and CFU counting. An additional 9 volumes of 2xYT supplemented with 20 mM glucose was added to the remaining culture and incubated overnight at 37 °C with shaking (225 rpm). The library was then extracted through midi-prep of the overnight culture.

### Selection of Sxkmer-loop variants

Selection cells (S2060 *ΔcadCBA* cells harboring plasmids pBL071 (cCadC-Aβ42) and pBL140 (P_*cadBA*_ cat)) were transformed by electroporation with the Sxkmer-loop library. Cells were allowed to recover for 1 h in 2xYT supplemented with 20 mM glucose, after which 200 ng/mL (to induce cCadC-Aβ42) and either 1 mM or 0.01 mM IPTG (to induce binders) was added to induce cCadC-Aβ42 and binder expression and the cells allowed to grow for another 2 h. A total of 10^8^ transformants were then plated on LB agar containing maintenance antibiotics, 200 ng/mL aTc and varying levels of chloramphenicol and IPTG. Plates were incubated at 37 °C for 1.5 days. Colonies were scraped and plasmids extracted by mini-prep. The Sxkmer-loop sequences were then amplified by PCR and cloned into the pBL098 backbone by USER assembly. Assembled plasmids were directly transformed into selection cells by electroporation. The same outgrowth procedure described above was repeated before cells were plated onto LB agar containing maintenance antibiotics, aTc, IPTG, and chloramphenicol. Surviving colonies were then analyzed by Sanger sequencing.

### Expression and purification of AβO binders

Overnight cultures of BL21 DE3 pLysS cells harboring expression plasmids of 6xHis-tagged Sxkmer variants were generated by inoculating single colonies into 4 mL of 2xYT containing maintenance antibiotics. Saturated overnight cultures grown overnight at 37 °C with shaking (225 rpm) were then diluted 1:100 into fresh 2xYT containing maintenance antibiotics in shake flasks and incubated at 37°C with shaking until the OD_600_ reached 0.4. Cells were then induced with the addition of 0.5 mM IPTG and grown for anotheer 4 h, at which point pellets were harvested by centrifugation and stored at -80°C until lysis.

Frozen cell pellets were thawed on ice and resuspended in lysis buffer (10 mM Tris-HCl, 0.1 mM EDTA, pH 7.5). Cells were lysed by sonication and the lysate cleared through centrifugation at 18,000 rcf for 1 h. Sxkmers were isolated using affinity chromatography with Nickel Sepharose resin (Cytiva). Purified protein was concentrated using Amicon Ultra-15 3000 Da molecular weight cutoff centrifugal filters and purified using size exclusion chromatography on a Superdex 75 10/300 column using the BioRad NGC Quest 10 Plus system. Pure fractions of Sxkmer variants were flash frozen in liquid nitrogen and stored at -80°C.

### Expression and purification of Aβ42 peptide

Purification of Aβ42 from inclusion bodies was performed as previously described.^67^ Briefly, overnight cultures of BL21 DE3 pLysS cells harboring an Aβ42 expression plasmid was diluted 1:100 into fresh LB media containing maintenance antibiotics and grown at 37°C with shaking (225 rpm) until OD_600_ of 0.8 was reached. 1 mL of this OD_600_ 0.8 culture was then transferred into 1 L of autoinduction media and incubated at 37°C for 14 h with shaking at 125 rpm. Cells were harvested by centrifugation and the pellet lysed by sonication (40% amplitude, [1 sec on, 1 sec off] for 1 minute total followed by 2 minutes incubation on ice, repeated for a total of 4 times) in 80 mL of lysis buffer (10 mM Tris-HCl, 0.1 mM EDTA, pH 7.5). The resulting lysate was spun down by centrifugation and the pellet re-suspended in 50 mL lysis buffer, then sonicated (40% amplitude, [1 sec on, 1 sec off] for 30 sec total). This step was repeated.

The lysate was spun down by centrifugation at 18,000 rcf for 8 minutes at 4 °C. The pellet was then dissolved in 50 mL of 8 M ice-cold urea and sonicated to dissolve the inclusion bodies (40% amplitude, [1 sec on, 1 sec off] for 1 minute total followed by 2 minutes incubation on ice, repeated for a total of 4 times). The resulting mixture was centrifuged at 18,000 rcf for 5 minutes at 4 °C. The supernatant was then added to 40 mL (settled resin volume) DEAE cellulose that was pre-equilibrated in ice-cold TE8 buffer (10 mM Tris-HCl, 0.1 mM EDTA, pH 8.5), and batch bound by incubating for 30 minutes on ice with gentle stirring with a glass rod. The bound resin was transferred to a Buchner funnel, washed with 200 mL of ice-cold TE8 buffer, then washed with 100 mL of ice-cold TE8 buffer containing 10 mM NaCl. Finally, the resin was eluted using 10 × 40 mL fractions of ice-cold TE8 buffer containing 50 mM NaCl. The eluates were flash frozen on dry ice and lyophilized.

Lyophilized fractions were combined and dissolved in ∼25 mL of room-temperature 6 M guanidine hydrochloride and purified by size exclusion chromatography using a Superdex 75 16/600 column. Collected fractions were flash-frozen on dry ice, lyophilized, and then re-dissolved in room-temperature 6 M guanidine hydrochloride and re-purified by SEC. Concentration was estimated using 280 nm absorbance of the center of the monomer peak assuming a molar absorptivity of 1440 (M^-1^ cm^-1^). Monomer fractions were portioned into 40 nmol aliquots, frozen on dry ice, lyophilized, and stored at -80°C until use.

### ThT fluorescence assays

A 500 mM stock of ThT (ChemCruz) solution was prepared in ThT buffer (20 mM NaPi, 0.2 mM EDTA, pH 8.0) by first sonicating to dissolve ThT then filtering through a 0.22 µm filter to remove particulates. Binder solutions at 2X the desired final concentrations were prepared in ThT buffer on ice in low protein binding microcentrifuge tubes. 40 µL of binder solution was transferred into a 96 well black-walled, half-area clear bottom, non-binding surface plate (Corning 3881).

Lyophilized Aβ42 aliquots were dissolved in 6 M guanidine hydrochloride and purified by SEC using a Superdex 75 10/300 column to obtain highly monomeric Aβ42. Aβ42 concentration was estimated using 280 nm absorbance of the center of the monomer peak assuming a molar absorptivity of 1440 (M^-1^ cm^-1^). A 2X master mix containing 6 µM freshly purified Aβ42 and 20 µM ThT was prepared in ThT buffer. 40 µL of the Aβ42/ThT master mix was added to the 40 µL binder solutions that had been prepared in the 96 well plate. The plate was immediately placed into a TECAN Infinite Mplex plate reader. The plate was incubated at 37 °C without shaking for 14-18 h with ThT fluorescence values read using a 440 nm excitation and 480 nm emission every 5 minutes. ThT fluorescence curves were analyzed using the AmyloFit software^60^.

### Transmission electron microscopy

Transmission electron microscopy (TEM) was performed at the University of Wisconsin School of Medicine and Public Health Electron Microscopy facility. Samples were prepared by incubating a 1:1 stoichiometry of 3 µM Aβ42 and binder for 4 h at 37 °C. 2 µL of sample was applied to a pioloform-coated 300-mesh thin Cu grid and stained using negative stain Nano-W (methylamine tungstate) (Nanoprobes) followed by water rinse. TEM images were captured using a FEI CM120 electron microscope with an BioSprint12 digital camera.

### Chemically stabilized AβO generation

Aβ42 oligomer species were generated as previously described^61,62^. Briefly, 1 mg HFIP treated Aβ42 peptide purchased from rPeptide was dissolved in HFIP (221 µL) and aliquoted into 10 × 0.1 mg aliquots. HFIP was evaporated under a gentle steam of nitrogen. For oligomer preparation, a single aliquot was dissolved in 100 µL 50 mM aqueous NaOH. This solution was then diluted into 767 µL PBS to a final concentration of 25 µM Aβ42 and incubated at 25 °C for 1 day to generate A11+ oligomers and 4 days for A11-oligomers.

### Dot blotting

2 µL of 25 µM Aβ42 monomer or oligomer preparations were spotted onto nitrocellulose membranes (Thermo Scientific) and allowed to dry for 5 min at room temperature. The membrane was then blocked by incubation in 3% BSA in Tris-buffered saline (TBS) pH 7.4 for 30 minutes at room temperature. This and all following incubations were performed with gentle agitation. Then, the membrane was incubated with 1 uM of Sxkmer-VKLVCVV in TBST (TBS + 0.1% TWEEN-20) at room temperature for 1 h. The membrane was washed by incubating in TBST for 10 minutes. This step was repeated for a total of 3 washes. The membrane was then incubated in a 1:1000 dilution of a mouse anti-His primary antibody (Invitrogen MA1-21315) for 1 h at room temperature. The TBST washes were repeated before the membrane was incubated in a 1:10000 dilution of HRP-conjugated anti-mouse IgG secondary antibody (Azure AC2115) for 1 h at room temperature. Finally, the membrane was washed three times with TBST. The membrane was treated with Radiance Q chemiluminescence substrate and imaged using an Azure c400 imager.

### Mass photometry

A 1 µM solution of Sxkmer-YRFFIYW was prepared in 20 mM NaPi, 0.2 mM EDTA, pH 8.0. Microscope cover slips were cleaned with alternating washes of H_2_O and isopropanol and dried under nitrogen. Sample well-containing silicon gaskets were added onto the coverslip and 3 µL of 1 µM Sxkmer-YRFFIYW was further diluted into 15 µL of 20 mM NaPi, 0.2 mM EDTA, pH 8.0 on the coverslip. 20 mM NaPi, 0.2 mM EDTA, pH 8.0 buffer alone was used as a control. Data was collected on a Refeyn OneMP mass photometer. After autofocus stabilization, 60 s movies were recorded, and the data was analyzed using the manufacturer’s software.

### Surface plasmon resonance

Aβ42 fibrils were prepared as previously described.^68^ Briefly, 30 µM Aβ42 solutions in 20 mM NaPi pH 8.0 buffer were incubated for 48 h at 37 C. The solution was then sonicated in a bath sonicator for 300 s at room temperature. This fibril-containing solution was then immobilized on a Cytiva Biacore SPR CM3 sensor chip using EDC-NHS amine coupling in a Biacore X. SPR was conducted with 100 µL injections at 10 µL/min flow rate of binder solutions in 20 mM NaPi 0.2 mM EDTA pH 8.0 buffer. Dissociation time was set at 330 s. Data was fit using single site binding model.

### Circular dichroism

Circular dichroism spectra were measured using 25 µM of binders in 20 mM NaPi 0.2 mM EDTA pH 8.0 buffer (400 µL in a 1 mm path length cuvette) on a Jasco J-1500 CD spectrometer at room temperature.

## Author Contributions

B.L. designed the research, performed the experiments, analyzed data, and wrote the manuscript. J.A.M. assisted with Aβ42 purification. T.W. designed and supervised the research and wrote the manuscript.

## Competing interests

The authors are in the process of filing a patent application on this work.

